# Multiple reproductive females in family groups of smooth-coated otters

**DOI:** 10.1101/2021.06.26.450053

**Authors:** Haaken Zhong Bungum, Mei-Mei Tan Heng Yee, Atul Borker, Chia Da Hsu, Philip Johns

## Abstract

Smooth-coated otters (*Lutrogale perspicillata*) are inhabitants of the waterways of India and Singapore. Otter families typically consist of a single mating pair with mature, nonbreeding siblings living in family groups, or “romps”. We note here the presence of multiple reproductive female otters within some romps, as well as the possible existence of simultaneous litters by different mothers. This phenomenon has not been recorded among *L. perspicillata* before. Here we address possible influences leading to multiple reproductive females within romps of smooth-coated otters, including inclusive fitness, incomplete suppression of reproduction, and existing in an urban environment. The numerous, recurring observations of multiple reproductive females warrant further investigation; while uncommon, this phenomenon is not as rare as once thought.

## Introduction

The evolution of communal and cooperative breeding systems are long-studied topics in animal behavior (e.g., Hayes 2000; Gilchrist 2006; Clutton-Brock 2009; Lukas and Clutton-Brock 2012a, 2012b; Federico et al. 2020), with various species relying on such systems to raise young and maintain their territories. Cooperative breeding systems are uncommon in mammals, present in about 5% of mammalian species (Lukas and Clutton-Brock 2012b). In these species, a group typically includes a single mating pair, while other members provide alloparental care for the offspring of the mating pair (Lukas and Clutton-Brock 2012b). Nonbreeding adults can be prevented from reproducing via suppressive means, such as infanticide or hormonal cues (Clutton-Brock 2009). The nonbreeders remain with the family into adulthood, often until either one or both of the breeding pair dies, are ousted, or the offspring disperse and form their own groups (Lukas and Clutton-Brock 2012b). In some species, such as giant otters (*Pteronura brasiliensis*), aging females are known to cease reproduction and become “grandmothers”, with younger females taking over the role of reproduction while the former matriarch provides parental care (Davenport 2010).

In contrast, communal breeding entails systems where multiple females in a group breed and share resources in raising young (e.g., Hayes 2000; Lukas and Clutton-Brock 2012b; Federico et al. 2020). Here reproductive suppression does not occur, and individuals can reproduce once sexually mature. Banded mongooses (Gilbert 2006) and numerous rodent species (Hayes 2000) are examples of communal breeders. Females in some rodent species may even share milk with non-offspring, i.e., allonurse them, which can be extremely costly (Hayes 2000). Benefits of communal breeding include improved thermoregulation (huddling), defense, and foraging (Hayes 2000).

While allonursing is common in communally breeding species (MacLeod et al. 2013), in cooperative systems where non-dominant females are generally not sexually active (Lukas and Clutton-Brock 2012b), allonursing can be especially costly to the nonbreeding individuals. Most potential benefits of allonursing among cooperative breeders seem to be kin-based, where dispersal is heavily delayed, and where groups consist mostly of close relatives (Lukas and Clutton-Brock 2012b, Federico et al. 2020). Among meerkats, allonursing females are often subordinate females who have recently been pregnant and return after having been evicted from the group (MacLeod et al. 2013).

### Multiple reproductive females in smooth-coated otters

Smooth-coated otters (*Lutrogale perspicillata*) are medium-sized predators (up to 10 kg) that are largely socially monogamous, living in family groups, or “romps”, that may include a dominant breeding pair (the “matriarch” and “patriarch”) and as many as 22 individuals from successive broods of offspring (personal observations). They perform several group-level behaviors, including group territorial defense, group defense against potential predators, and group foraging (personal observations). They can be the apex predator in many ecosystems. Smooth-coated otters live in Asia, especially South and Southeast Asia, often inhabiting mangroves, coastal rivers, and other waterways, including ones in Western India and Singapore.

Here we collate observations of romps of smooth-coated otters with what appear to be multiple reproductive females (**MRF**). We include data from camera traps, from our own personal observations, from interviews with other otter watchers, and gleaned from social media. Although social media is comprised of largely *ad libitum* observations (Altmann 1974), it is useful for recording uncommon events (Nelson and Fijn 2013). We include observations from rural sites in Goa, India, and from Singapore, one of the most densely populated countries in the world. Smooth-coated otters returned to Singapore after decades of absence (Khoo and Lee 2020). Singapore now has at least eleven distinct romps living in the city’s highly canalized waterways (Khoo and Sivasothi 2018). Some families are extremely habituated to humans, and a loose network of photographers, naturalists, students, and other scholars watch some romps almost daily. Territories are fairly stable, week to week, and observers can distinguish most families based on their locations, the number of animals in a group, and individual markings on some otters. We also include evidence from a necropsy of a dead otter pup found in Singapore.

We consider any of the following criteria to indicate multiple females exhibiting some reproductive activity within a romp: 1) pups of substantially different sizes and stages of development, indicating they are not from the same litter; 2) more than one female in a romp with enlarged nipples; 3) more than one female in a romp actively nursing pups. We consider the presence of females with enlarged nipples as likely evidence of recent birth or lactation.

Although smooth-coated otters primarily practice cooperative breeding (Hussain 1996; Hwang and Larivière 2005; Sivasothi and Khoo 2018b), we note eight instances of romps with MRF between 2015 and 2021: two in Goa and six in Singapore, with one Singaporean romp having had three MRF events over four years (**Table 1**).

**Table 1.**
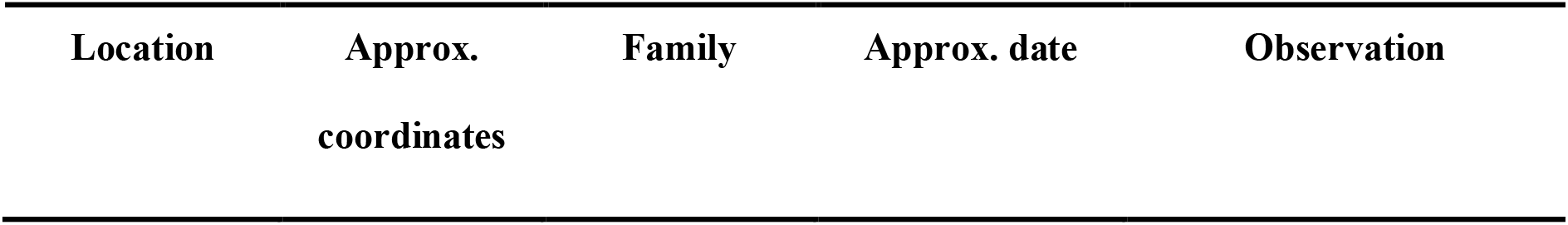
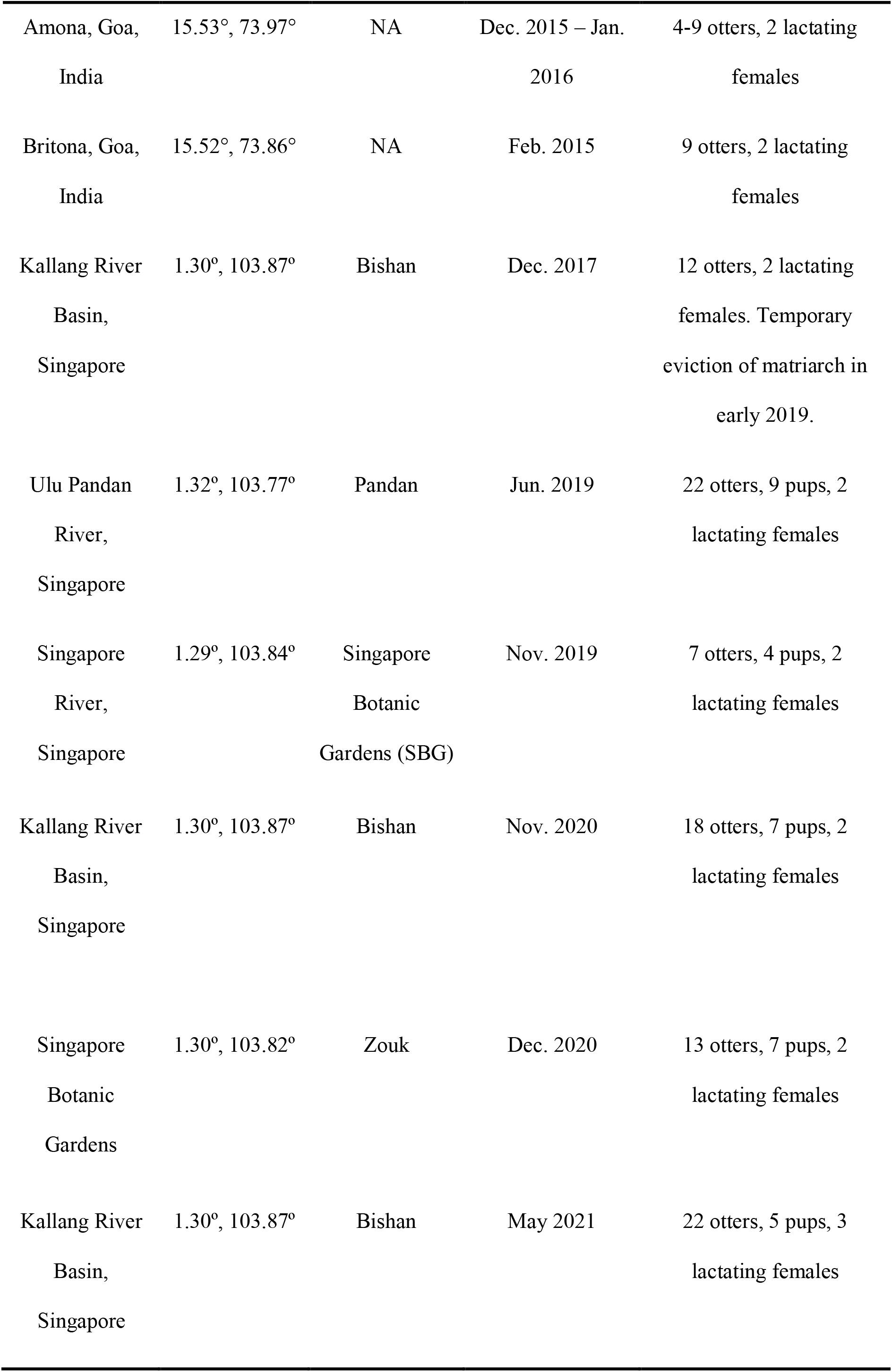
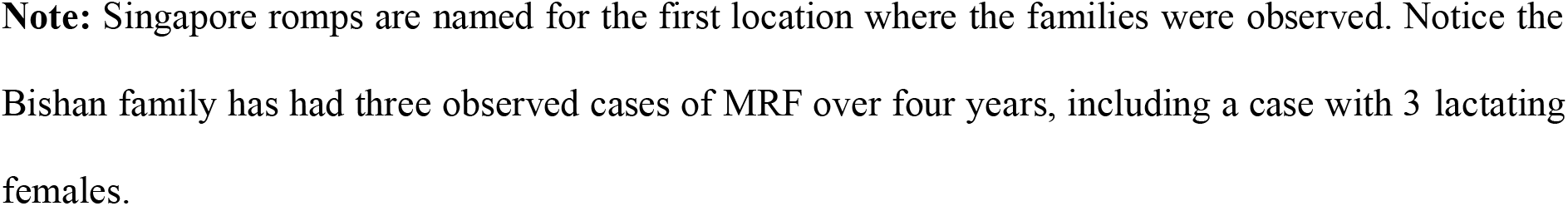
Observations of romps with multiple reproductive female smooth-coated otters in Goa, India, and Singapore.

Members of the research organization Wild Otters placed video camera traps at several otter latrine (“spraint”) sites, including in Amona and at an otter holt in Britona (**Table 1**). Both locations include mangrove habitats with abundant prey for smooth-coated otters. Both observations of MRF in India entailed short videos from these camera traps. In each case a romp of otters included two different females that had substantially enlarged nipples (e.g., Fig 1E).

**Figure 1.**
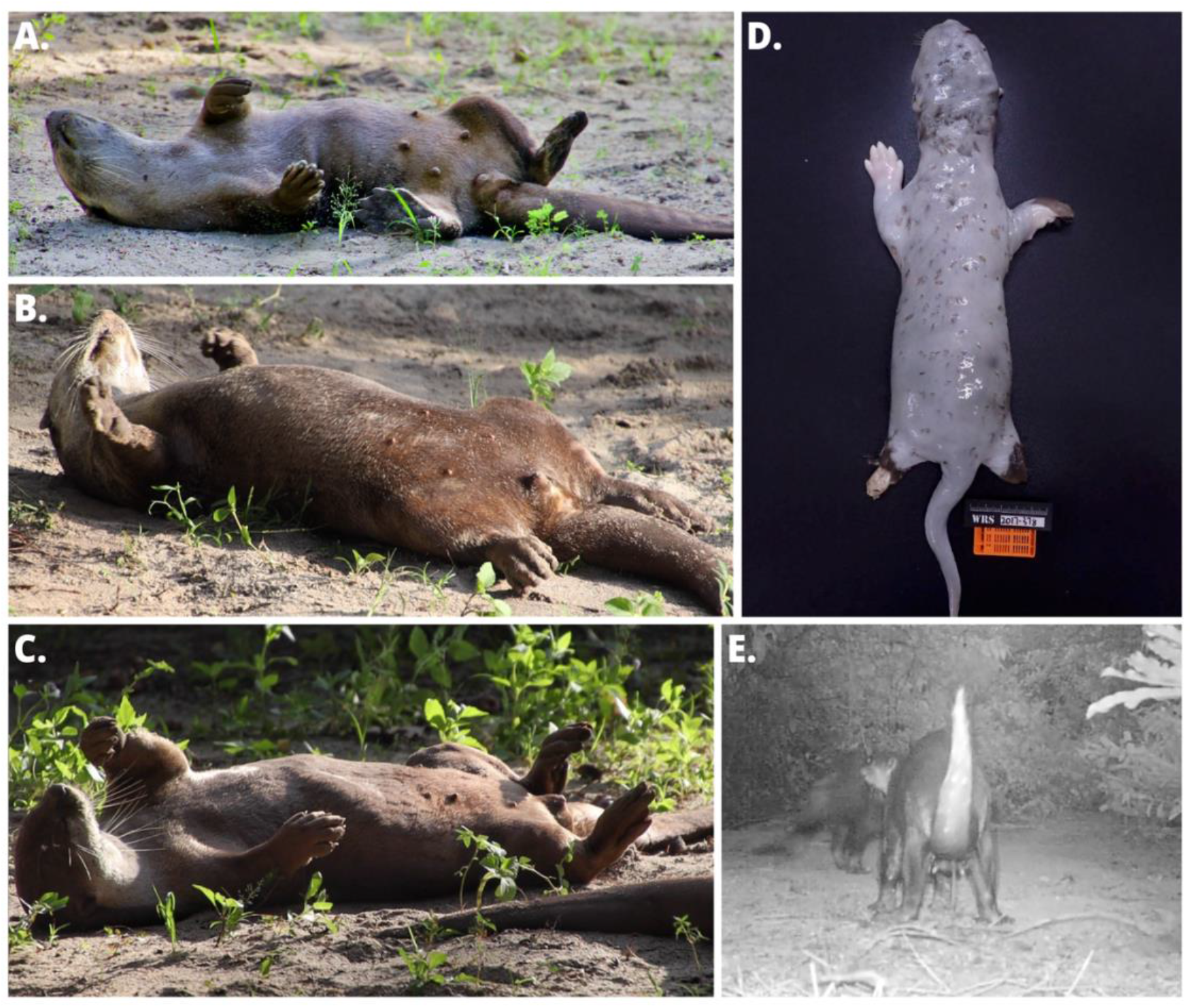
Examples of confirmed multiple-lactating-females from the Bishan family, June 2021. A-C were taken on the same day, showing the three mothers within the romp; C is the matriarch, identified by her 5^th^ nipple. D was taken from the necropsy of the dead pup from 2017, a suspected infanticide (see text). E was taken in Amona, India, in early 2016; two females were lactating in the romp.

The earliest observation of MRF in Singapore occurred about two weeks after the Bishan matriarch’s 4th litter emerged in late 2017 (Khoo and Sivasothi 2018). Otter watchers spotted a subordinate female (“Crazy Sis”) of the Bishan family carrying a small pup. The emerging pups were much larger and probably weeks older; smooth-coated otter pups typically emerge from their holts around six weeks after birth and start to swim about a week later (Khoo and Sivasothi 2018). Days later, a small pup was found dead near the Bishan holt. A necropsy concluded the cause of death was multiple bite wounds from conspecifics (Fig 1D). Although the necropsy suggests territorial conflict as the cause (Supp. Fig 1), and the Bishan family had been involved in a series of intense and lethal territorial disputes months prior (personal observations), we know of no territorial conflict between Bishan and other romps at the time of the pup’s death. A possible cause of death is wounds inflicted by adults in the Bishan romp. If so, this is the only recorded instance of infanticide in smooth-coated otters by family members.

In May 2019, the Pandan romp consisted of nine adults and nine pups, an unusually large number of pups, given a previously recorded mean litter size of 4.86 (Khoo and Sivasothi 2018); these pups likely belonged to litters from two females. June 2019 photographs confirmed the existence of two MRF (Table 1; Supp. Table 1), although there was no indication of which female each pup belonged to.

The Singapore Botanic Garden (SBG) family originally consisted of two same-aged sisters from the Marina family and a male of unknown origins. In November 2019, one sister gave birth to one pup, then the second sister (“Lightbulb”) gave birth to three more (Table 1; Supp. Table 1). This romp has since suffered several mortalities, and Lightbulb disappeared in 2020. The matriarch now has a new partner, giving birth to four pups in April 2021.

In late 2020, the Bishan matriarch had her seventh litter. Weeks after her pups emerged, a much smaller pup was spotted with one of the subordinate adult females in the romp (Table 1). The family abandoned the pup, which had yet to open its eyes when it was euthanized by the authorities. There is no evidence of infanticide (Supp. Table 1). The mother of this pup gave birth to two pups later in March 2021, along with 3 from another subordinate female at the same time (Table 1). The matriarch and both subordinate females have been observed nursing these 5 new pups, (e.g., Fig1A-C; File S1) and in May 2021 the romp had 22 members (Supp. Table 1).

In December 2020, the Zouk romp displayed MRF, with the matriarch and a daughter (“Flowerhead”) each having enlarged nipples (Table 1). Although it is uncertain whether the group of seven Zouk pups were from one or two mothers, pups were observed nursing from both females (Supp. Table 1)

## Discussion

The repeated occurrence of MRF in India and Singapore suggests that these events are uncommon but not especially rare. We review several hypotheses, including ones related to incomplete suppression of reproduction, inclusive fitness, and urban environments (**Table 2**).

**Table 2.**
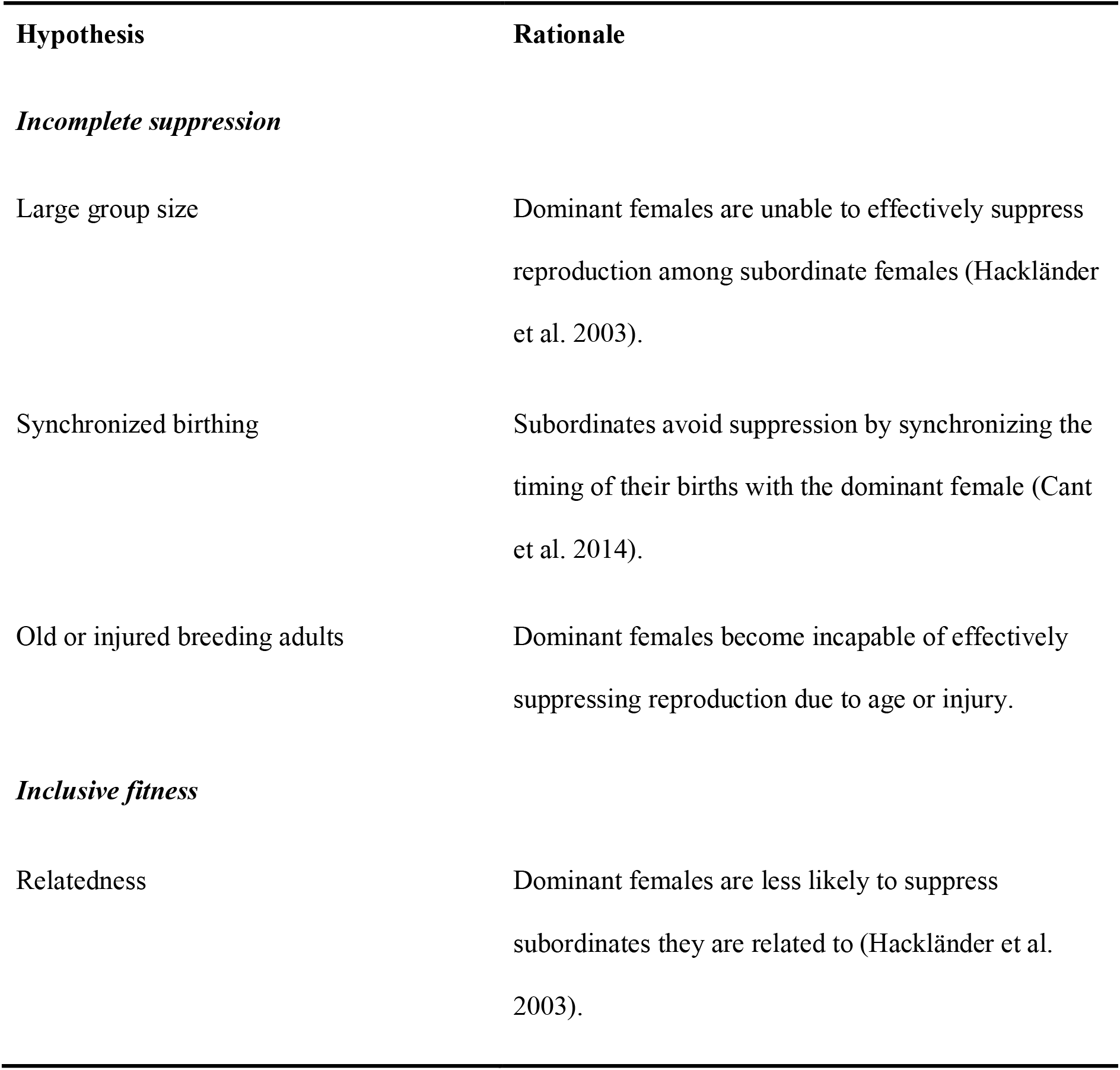
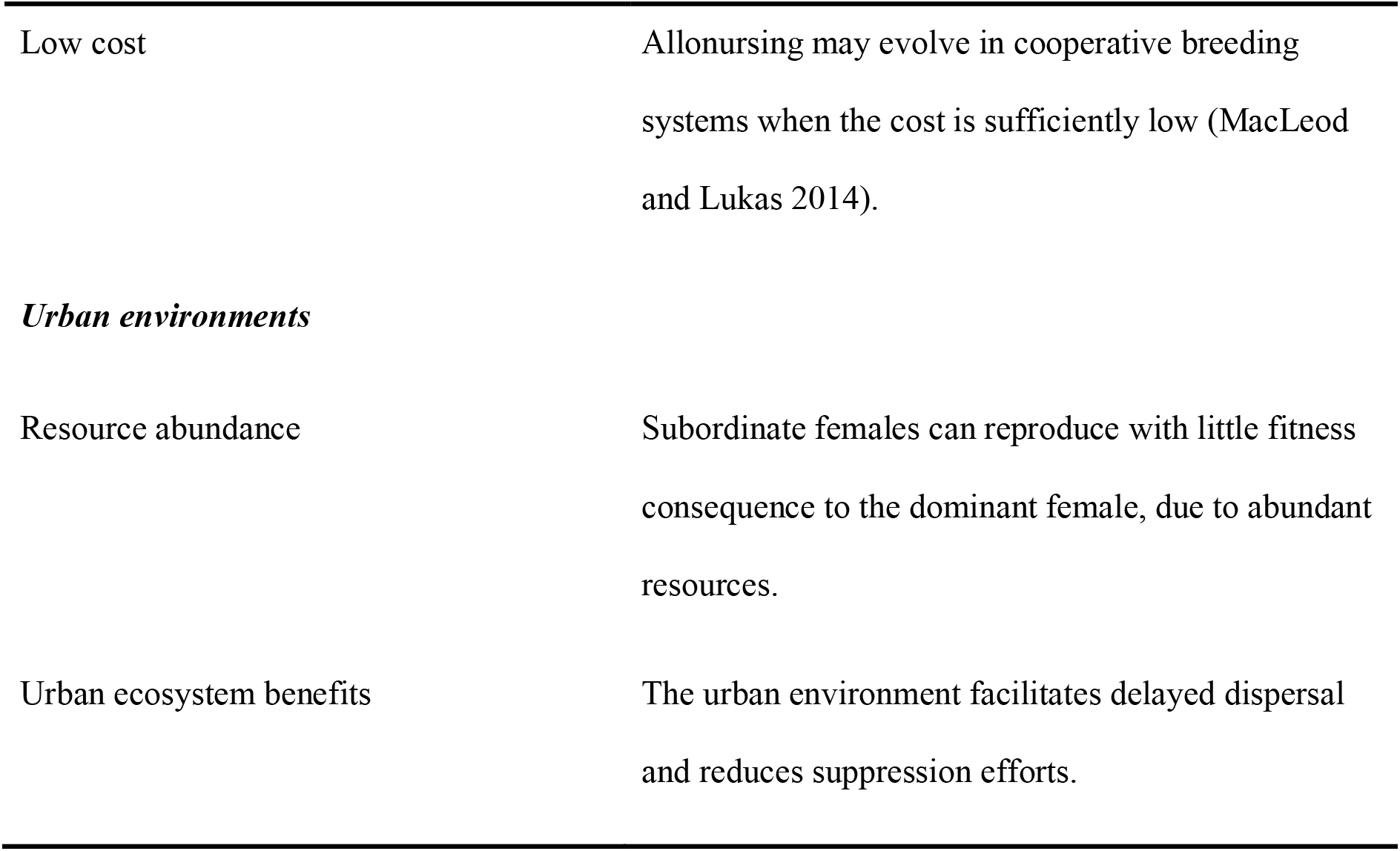
Hypotheses for the cause of multiple reproductive otter females

| Hypothesis | Rationale |
| --- | --- |
| *Incomplete suppression* |  |
| Large group size | Dominant females are unable to effectively suppress reproduction among subordinate females (Hackländer et al. 2003). |
| Synchronized birthing | Subordinates avoid suppression by synchronizing the timing of their births with the dominant female (Cant et al. 2014). |
| Old or injured breeding adults | Dominant females become incapable of effectively suppressing reproduction due to age or injury. |
| *Inclusive fitness* |  |
| Relatedness | Dominant females are less likely to suppress subordinates they are related to (Hackländer et al. 2003). |
| Low cost | Allonursing may evolve in cooperative breeding systems when the cost is sufficiently low (MacLeod and Lukas 2014). |
| ***Urban environments*** |  |
| Resource abundance | Subordinate females can reproduce with little fitness consequence to the dominant female, due to abundant resources. |
| Urban ecosystem benefits | The urban environment facilitates delayed dispersal and reduces suppression efforts. |

### Hypotheses for multiple reproductive females

The first evolutionary step towards communal or cooperative breeding may be delayed dispersal, and the distinction separating the two breeding systems is reproductive suppression (Federico et al. 2020). When successful, suppression enables the dominant mating pair to monopolize breeding and to recruit other individuals to provide various levels of care for the offspring. Suppression can occur via several means, such as hormonal cues, aggression and physical violence towards subordinates, and infanticide (Spiering et al. 2010; Lukas and Huchard 2019). Suppression is present in giant (*Pteronura brasiliensis*) and small-clawed (*Aonyx cinereus*) otters (Groenendijk et al. 2014; Perdue et al. 2013), which are also cooperative breeders. Notably, MRF has been recorded in giant otters, although this is limited to one observation (Leuchtenberger and Mourao 2009).

In marmots, the frequency and severity of agonistic suppression varies depending on the relatedness of the matriarch to other females in the group (Hackländer et al. 2003). Furthermore, as group size increases, dominant females are less fertile, suggesting there is a cost associated with policing behavior (Hackländer et al. 2003). Dominant female banded mongooses also police subordinate female reproduction. Subordinate females appear to synchronize births to increase pup survival rates; dominant females are less likely or less capable of committing infanticide, leading to higher survivorship, while subordinate females’ pups born before the dominant female’s pups are more often killed (Cant et al. 2014). This pattern suggests that dominant females are not able to effectively distinguish their own offspring from others when they are of similar age.

We report evidence of what may be policing by the Bishan dominant female of a subordinate female’s pup (Fig 1D). Smooth-coated otters sometimes violently evict other otters from romps. After several other social changes to the Bishan family, including the death of the Bishan patriarch and the subsequent takeover of the romp by another adult male (“Scarface”), a Bishan daughter (“Crazy Sister”) evicted the Bishan matriarch from the romp, before apparently being evicted herself by another subordinate (“White Tip”). These conflicts, which occurred between April and July 2019, ended with all three females rejoining the romp at different times, remaining until present (Supp. Table 1). Such conflict suggests a dynamic like that of mongooses (MacLeod et al. 2013), where allonursing may follow pregnancy, eviction, and return, of subordinate females. We suspect policing may also have occurred with the birth of the small Bishan pup, although there was no actual physical injury, and the subordinate female abandoned the pup without any known conflict.

Large smooth-coated otter romps may consist of two, three, or more, successive broods of offspring, and the dominant pair may be several years old. Therefore, one possibility for MRF is that the matriarch is simply too old or the romp too large to effectively suppress reproduction. Two of the Singapore otter families (Bishan, Pandan) were among the largest, each with 22 members at their peaks. However, small, young romps such as SBG have been observed with MRF, and the Indian romps experiencing MRF were not especially large, suggesting aging matriarchs or overly large romps are unlikely to be the only cause of MRF. In SBG’s case, the subordinate female even had more pups (three) than the presumed dominant female (one). Other factors may contribute to MRF, perhaps including stress and injury associated with violent group territorial conflict between rival romps. However, while such “otter wars” do occur between romps of Singapore’s otters, these territorial disputes have not been observed in all cases of MRF. Violent territorial defense may contribute to MRF but it too is unlikely to be an only cause.

Allonursing between mothers and daughters does have potential inclusive fitness benefits, and smooth-coated otter romps typically consist of a breeding pair and their offspring, which are likely to be full siblings. Therefore, these inclusive fitness benefits may be substantial, and they contribute to allonursing in other systems (e.g., Hackländer et al. 2003). In free-ranging dogs, pups have even been observed to steal milk from other mothers (Paul and Bhadra 2017). The cost of allolactation may not be especially high depending on the conditions (MacLeod and Lukas 2014), which could favor allolactation in smooth-coated otters. However, the evidence of policing and the observations of females’ eviction suggest potential within-group conflict related to reproduction. How these individual and inclusive fitness benefits interact is not clear.

Cooperative breeding has been intensely scrutinized, and while the traditional explanation is that kinship plays a large role in obtaining these indirect fitness benefits, another body of evidence suggests that group dynamics among non-kin could also contribute to the development of this breeding system (Lukas and Clutton-Brock 2012b). Groups of cooperative breeding giant otters include unrelated individuals, for example (Ribas et al. 2015), suggesting that members of different giant otter families sometimes merge to form romps. At least some of Singapore’s smooth-coated otter romps are closely enough watched that merging of unrelated families into one romp seems unlikely; otter watchers can trace the growth of romps brood by brood. But without a more detailed analysis we cannot know how kinship and other group dynamics contribute to the development of MRF within smooth-coated otters.

Another factor influencing MRF may be smooth-coated otters’ existence in urban Singapore. In other animals that returned to Singapore recently, some notable behaviors have been attributed to the urban environment. For example, pied hornbills have returned to Singapore in the last two decades, and they are frequently observed eating nestlings and fledglings of other birds, perhaps because those nestlings are easy to find in an urban environment (Loong et al. 2021). Conditions unique to an urban environment may also favor otter romps with multiple reproductive females.

The overall population density of smooth-coated otters in Singapore seems to be high. This high density suggests that they may enjoy unnaturally favorable conditions. Indeed, many of Singapore’s waterways are full of introduced fish. By one estimate, Singapore’s otters prey on large, introduced species of cichlids more than 90% of time while in Singapore’s freshwater reservoirs (Theng et al. 2016). Conditions with ample prey and few predators could potentially favor delayed dispersal and impose few fitness costs on the dominant female if subordinate females reproduce. Alternatively, abundant resources could lead to fewer ecological constraints and greater fitness gains if subordinate animals disperse early and establish their own territories. This latter hypothesis could lead to smaller territories among otter clans than in less abundant environments. The interplay between ecological constraints, demographics of a long-lived carnivore, and inclusive fitness is complex (e.g., Emlen 1982, Hatchwell and Komdeur 2000, Pen and Weissing 2000, Frederico et al. 2020); our aim is not to disentangle those factors here. However, we observed MRF in both urban Singaporean environments and rural Indian environments. Therefore, we posit that it is unlikely MRF is solely a consequence of urban ecological factors, such as abundant forage, or absence of predators.

## Conclusion

We note several occurrences of multiple reproductive females within smooth-coated otter romps, eviction of subordinate females, and what may be infanticide by romp members. The growing pool of observations suggest that MRF in smooth-coated otter romps is not especially rare. Its occurrence in several families in Singapore, including repeated instances in one romp, suggests that this largely cooperatively breeding mammal has a social system more complex than originally thought. We consider hypotheses that may contribute to MRF but find that none by itself adequately explains MRF in smooth-coated otters.

## Acknowledgments

We would like to thank the otter watching communities of Singapore for their helpful comments and conversations, especially Patrick Ng, Jeff Teo, Jeff Tan, and the contributors to several Singapore-based Facebook pages, including Ottercity, OtterWatch, Omni Channel, Otter Channel, Myottermelon, and the YouTube channel RandomSG. We would like to thank Hannah Krupa, Abhishek Gopal, and Komal Gogi, for their field work in Goa, India. We would also like to thank Nicole Duplaix for her helpful input on earlier drafts of this manuscript. This study was conducted in partial fulfilment of the Yale-NUS College Capstone requirements for Mei-Mei Tan and in partial fulfilment of the Yale-NUS College Independent Research requirements for Haaken Bungum.

## Supplementary materials

**Table S1.**
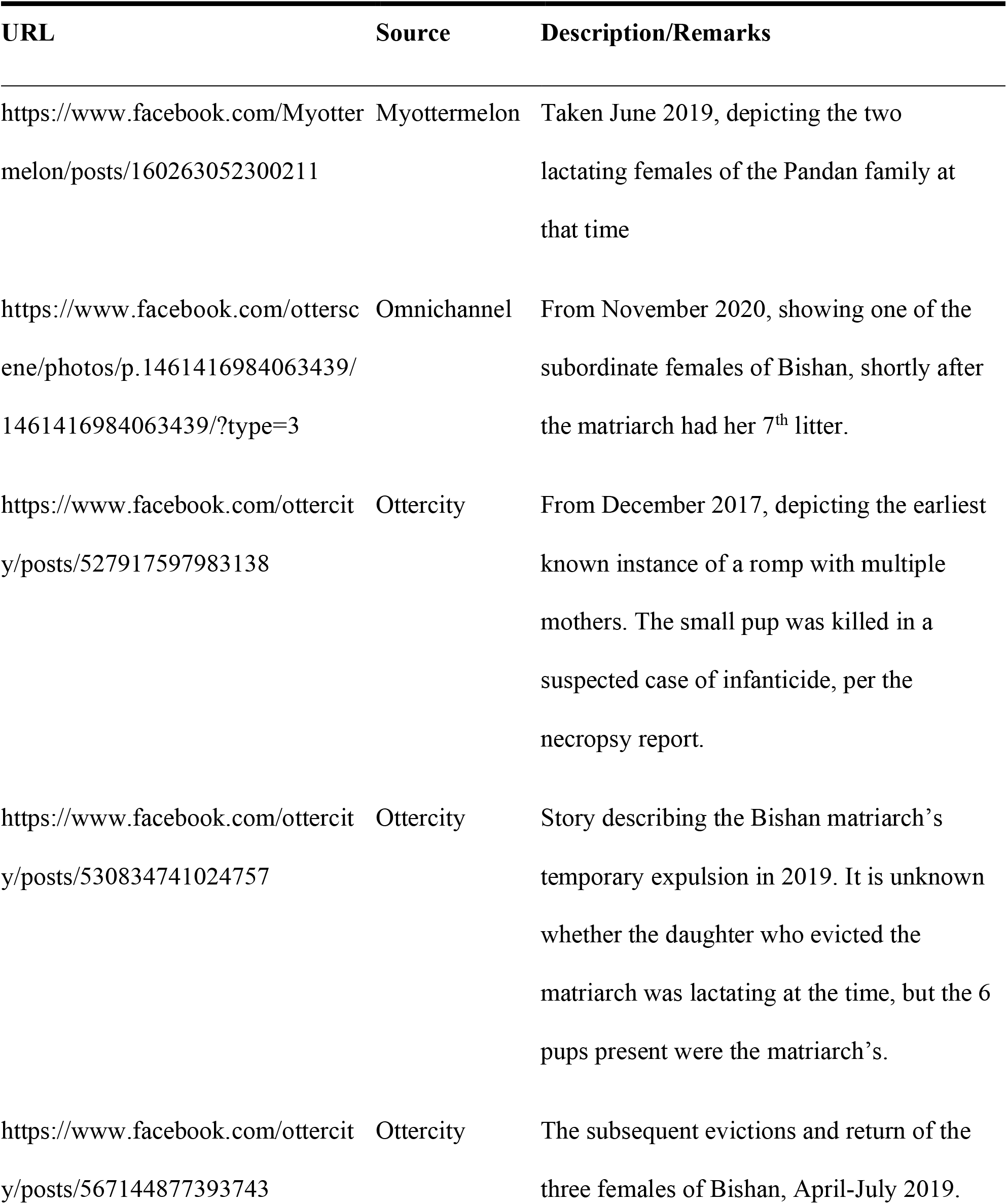
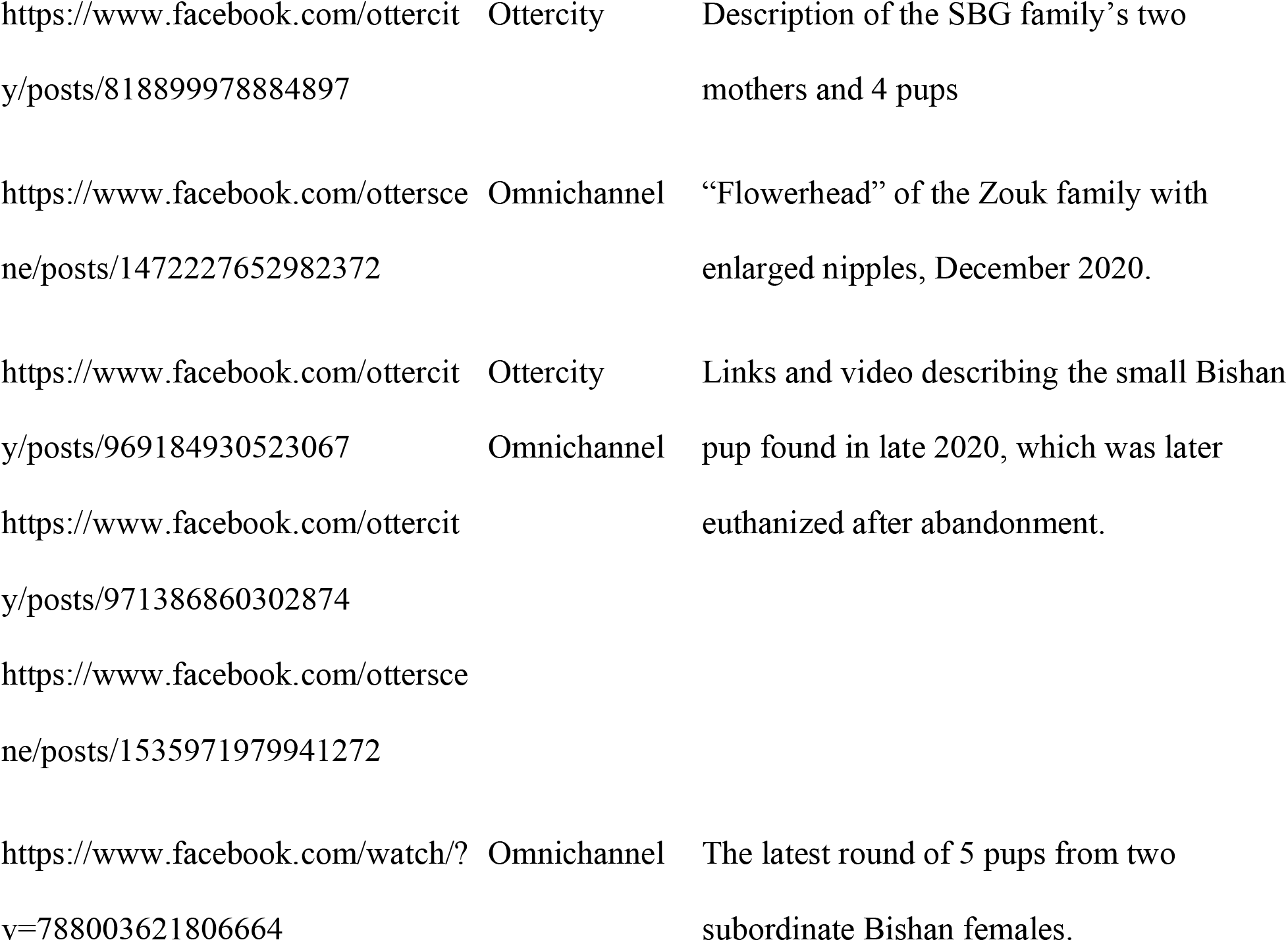
Links to relevant photographs, videos, and chronicles, from public otter-watching groups on Facebook.

**File S1**. Video of Bishan romp “grooming” in which the matriarch and two subordinate females with enlarged nipples are visible.

**Figure S1.**
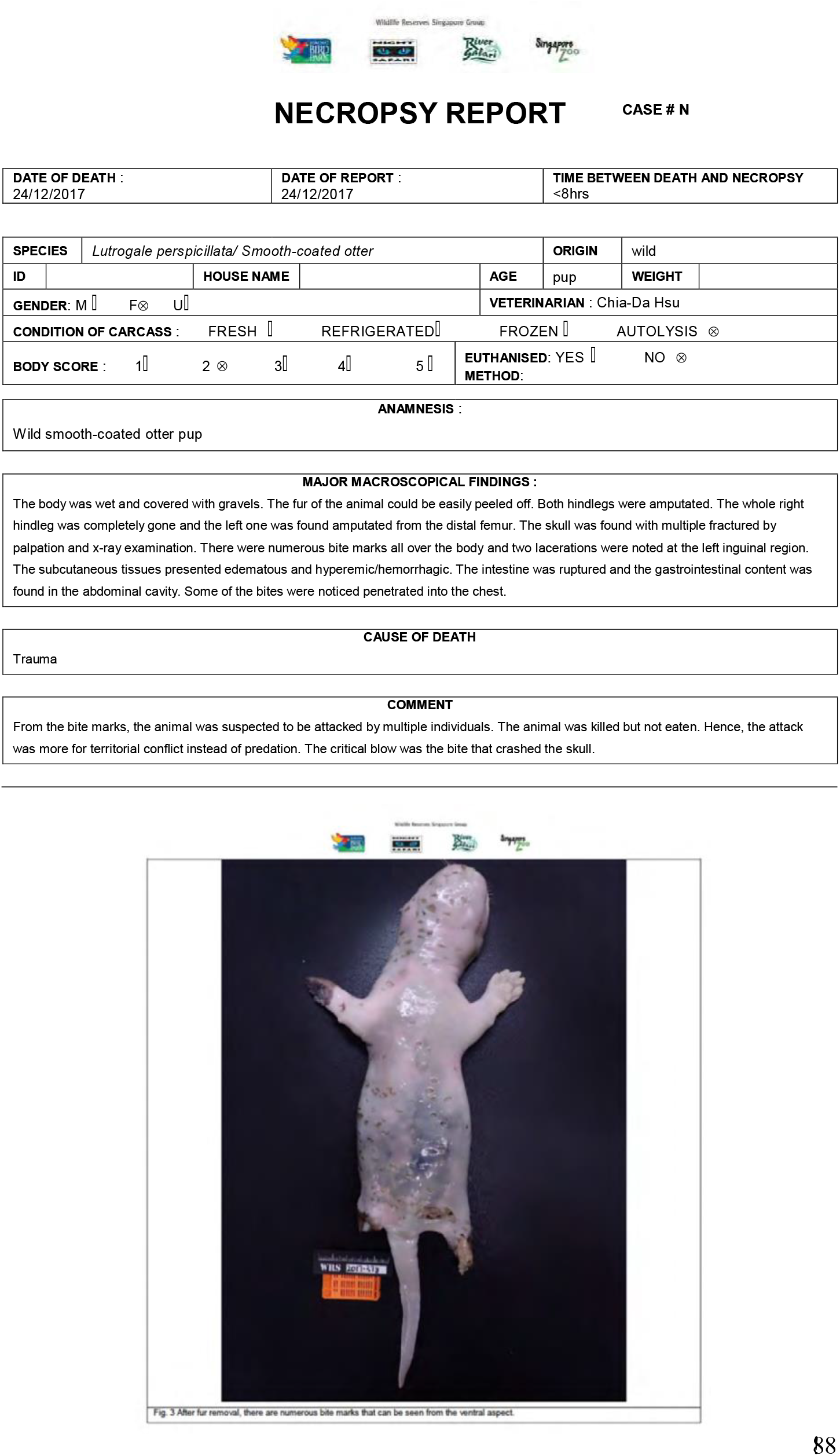
Necropsy report of dead otter pup found near Bishan holt.

## Notes

### Competing Interest Statement

The authors have declared no competing interest.

